# Benchmarking state-of-the-art approaches for norovirus genome assembly in metagenome sample

**DOI:** 10.1101/2022.07.05.498785

**Authors:** Dmitry Meleshko, Anton Korobeynikov

## Abstract

**Motivation:** A recently published article in BMC Genomics by Fuentes-Trillo et al (2021) contains a comparison of assembly approaches of several Noroviral samples via different tools and preprocessing strategies. Unfortunately the study used outdated versions of tools as well as tools that were not designed for the viral assembly task. In order to improve the suboptimal assemblies the authors suggested different sophisticated preprocessing strategies that seem to make only minor contributions to the results. We redone the analysis using state-of-the art tools designed for viral assembly.

**Results:** Here we demonstrate that tools from the SPAdes toolkit (rnaviralSPAdes and coronaSPAdes) allows one to assemble the samples from the original study into a single contig without any additional preprocessing.

## Background

Novel virus discovery is a very popular topic in bioinformatics nowadays including large-scale studies (Edgar et al, 2022; Kawasaki et al, 2021). Many research groups switched their attention to viral studies therefore exploding the amount of papers devoted to the subject. Despite sufficient interest to this topic, there is no established state-of-the-art method to perform a viral genome assembly. This fact can be explained by the overall diversity of the viral genomes and the corresponding sequencing data: one cannot expect that viral genome assembly approaches suitable for dsDNA bacteriophages can be extended without modifications to RNA viruses. As a result, some papers are naturally devoted to benchmarking of viral assembly approaches for different kinds of input data. However, proper benchmarking design (cf. (Luo et al, 2009; Magoc et al, 2013; Meyer et al, 2022; Sczyrba et al, 2017)) and interpretation of obtained results is a non-trivial task and flaws made here could easily lead to somewhat controversial results.

Recently, an article by Fuentes-Trillo et al (2021) discussing various approaches for Norovirus genome assembly was published in BMC Genomics. Using the approaches presented in this article we want to highlight some common benchmarking problems that might result in misleading conclusions and that can be easily avoided.

Firstly, the article was submitted to the journal and subsequently published in the year 2021, but the versions of tools used are extremely outdated. In particular, the authors compared metaSPAdes v.3.11.1 (Nurk et al, 2017) and MEGAHIT v.1.1.3 (Li et al, 2015) genome assemblers. metaSPAdes 3.11.1 was released back in March 2018, while the current version of SPAdes is 3.15.4 and SPAdes team makes multiple releases each year. Same problem does exist with MEGAHIT since v.1.1.3 was also released in March 2018, while the newest version is 1.2.9 that was released later in October 2019. Use of outdated tools together with claims that one tool “performed better” might be misleading and do not necessary reflect the current situation. This is especially important for benchmarking studies (as compared to papers that provide novel biological insight), since such comparison is the main result of the paper.

Secondly, the authors for some unknown reason have chosen tools that are not suitable for viral assembly problem and no further justification were made on such choice. We note that metaSPAdes and MEGAHIT are metagenomic assemblers. Though both assemblers have proven themselves even in non-metagenomic settings including viral assemblies (Roux et al, 2017; Sutton et al, 2019), the fact that authors did not try specialized assemblers that work with RNA and RNA viral data is worrying and might serve as an example of improper benchmark design. Indeed, Noroviruses are RNA viruses and surprisingly no single transcriptome or viral assembler was evaluated. Even in 2018 there were multiple prominent assemblers to try including e.g. Trinity (Grabherr et al, 2011), rnaSPAdes (Bushmanova et al, 2019), Savage (Baaijens et al, 2017), IVA (Hunt et al, 2015)) among the others. Also recently a dedicated RNA viral assembler rnaviralSPAdes (Meleshko et al, 2022) was developed (the preprint and the tool itself were available from summer 2020).

Finally, the presentation of several statistics that authors used to show the results could be improved. Authors assembled 8 datasets and reported multiple mean values across these datasets. These values might vary a lot during assembly and taking mean N50, contig size, number of contigs provides very limited information about the assembly results. As an immediate outcome, the benchmarked approaches can hardly be compared directly basing on these values. Although the authors aligned contigs using BLAST and reported mean values here are slightly more informative, we need to note that that there are only 8 samples. Therefore the resulting tables could possibly be reformatted so the reader can compare various statics across the samples and assess their variability.

Unfortunately, these inaccuracies were somehow slipped through the peer review process. We decided to try and reproduce the analysis from the mentioned study using the state of the art version of tools and show that the majority of them do correctly assemble noroviral datasets in question into a single contig without any additional sophisticated clustering procedures (that were presented in the article as a way to overcome assembly deficiencies). Luckily, benchmarks performed by the authors are easily reproducible, all data is available and versions of tools used are stated.

## Results

Scaffolds produced by each assembler were aligned to corresponding references using QUAST. The results obtained are summarized in Table 1. They clearly show that rnaviralSPAdes and coronaSPAdes (version of rnaviralSPAdes that could use profile HMM models to guide an assembly) are better suited for assembly of this data than rnaSPAdes as well as metaSPAdes and MEGAHIT (these two were benchmarked by Fuentes-Trillo et al (2021)). We note that each sample was assembled into a single contig with perfect or near-perfect quality.

**Table 1.** Benchmarking of assemblers rnaviralSPAdes (RVS), coronaSPAdes (CS), rnaSPAdes (RS), MEGAHIT (M) and Trinity (T) on several Noroviral datasets. Best results are outlined in bold.

|  |  | RVS | CS | RS | M | T |
| --- | --- | --- | --- | --- | --- | --- |
| SRR8074276 | Longest alignment (nt) | 7,538 | 7,538 | 7,538 | **7,548** | **7,547** |
|  | Genome fraction% | 99.83 | 99.83 | 99.83 | **99.96** | **99.94** |
|  | Longest alignment IDY% | **99.95** | **99.95** | 99.91 | 99.87 | 99.93 |
| SRR9141472 | Longest alignment (nt) | **7,569** | **7,569** | 5,282 | 7,560 | 5,848 |
|  | Genome fraction% | **100.0** | **100.0** | 78.80 | 99.88 | **100.0** |
|  | Longest alignment IDY% | **100.0** | **100.0** | 99.95 | 99.99 | **100.0** |
| SRR9141473 | Longest alignment (nt) | **7,536** | **7,536** | 4,979 | 7,487 | 6,899 |
|  | Genome fraction% | **100.0** | **100.0** | 90.55 | 99.35 | **100.0** |
|  | Longest alignment IDY% | **100.0** | **100.0** | 99.91 | 99.97 | **100.0** |
| SRR9141474 | Longest alignment (nt) | **7,541** | **7,541** | 6,838 | 7,516 | **7,542** |
|  | Genome fraction% | **99.99** | **99.99** | 99.99 | 99.65 | **100.0** |
|  | Longest alignment IDY% | **100.0** | **100.0** | 99.91 | 99.99 | **100.0** |
| SRR9141475 | Longest alignment (nt) | 7,482 | 7,493 | 7,049 | 7,440 | **7,534** |
|  | Genome fraction% | 99.28 | 99.43 | 99.08 | 98.72 | **99.97** |
|  | Longest alignment IDY% | **100.0** | **100.0** | 99.96 | **100.0** | **100.0** |
| SRR9141476 | Longest alignment (nt) | **7,533** | **7,533** | 5,699 | 7,526 | **7,533** |
|  | Genome fraction% | **100.0** | **100.0** | **100.0** | 99.90 | **100.0** |
|  | Longest alignment IDY% | **100.0** | **100.0** | 99.98 | **100.0** | 99.98 |
| SRR9141477 | Longest alignment (nt) | **7,554** | **7,554** | 5,935 | 7,467 | 3,658 |
|  | Genome fraction% | **99.95** | **99.95** | 91.20 | 98.79 | 99.84 |
|  | Longest alignment IDY% | **100.0** | **100.0** | 99.93 | 99.96 | 99.97 |
| SRR9141478 | Longest alignment (nt) | **7,540** | **7,540** | 6,592 | **7,540** | **7,540** |
|  | Genome fraction% | **100.0** | **100.0** | 95.90 | **100.0** | **100.0** |
|  | Longest alignment IDY% | **99.99** | **99.99** | 99.95 | **99.99** | 99.88 |

Moreover, the results clearly show that one do not need any additional pre- and post-processing steps beyond quality trimming to obtain good results contrasting with the assembly pipeline presented in (Fuentes-Trillo et al, 2021) that included multiple steps of read binning, contamination filtering and norovirus read filtering. All these steps might influence the final result and cause the degradation of assembly quality in general.

Since our approach only includes read trimming as a preprocessing step and therefore can be directly compared to pC approach of (Fuentes-Trillo et al, 2021). Here we see that coronaSPAdes was able to assemble 7 out of 8 samples into a contig longer than 7,500 bp, and the contig length of the remaining sample is 7,493 bp that places this sample into a near-complete category. Original pC approach utilized metaSPAdes and MEGAHIT, which assembled 4 out 8 and 5 out of 8 samples into a contig longer than 7,500 bp correspondingly.

## Discussion

Genome assembly task is a very hard but well studied computational problem. Multiple genome assemblers are available, and the choice of the assembler is highly dependent on input data properties and even the result desired. Nevertheless the choice of an assembler suitable for a given kind of input data cannot guarantee a complete genome assembly for complex datasets. However we emphasize that even for datasets with low complexity the researcher should choose an assembler carefully. A common problem seen in papers of “benchmarking” kind is the usage of improper tools or their obsolete versions.

We showed that specialized RNA viral assemblers such as coronaSPAdes and rnaviralSPAdes were able to outperform metagenomic assemblers metaSPAdes and MEGAHIT and RNA-assemblers Trinity and rnaSPAdes on the noroviral assembly task.

## Conclusion

Our experiments showed that the genome assembly using specialized tools and latest version of these tools yields better results in terms of correctness and contiguity. Ad-hoc assembly approaches ended in worse results. Moreover, the labor costs associated with such approaches are much higher because suboptimal results force researchers to find a way to improve initial results, that is usually harder than assembly itself. Finally, this short article emphasize the importance of specialized tool development and promotion.

## Methods

Raw data was trimmed using BBDuk as in (Meleshko et al, 2022) and assembled using coronaSPAdes 3.15.4, rnaviralSPAdes 3.15.4, rnaSPAdes 3.15.4, Trinity 2.13.2 and MEGAHIT 1.2.9 in default mode. Norovirus HMM models for coronaSPAdes were extracted from RVDB-prot-HMM database (Bigot et al (2020)).

### Data availability

Noroviral HMMs that were used for coronaSPAdes assembly are available at https://cab.spbu.ru/software/coronaspades/.

### Author’s contributions

Both authors conceived, designed, performed the analysis and wrote the paper.

## Acknowledgements

The research was carried out in part by computational resources provided by the Resource Center “Computer Center of SPbU”. The authors are grateful to Saint Petersburg State University for the overall support of this work. The authors were supported by the Russian Science Foundation (grant 19-14-00172).

