## Supplementary Materials for "Benchmarking state-of-the-art approaches for norovirus genome assembly in metagenome sample"

July 5, 2022

### 1 Software versions

1. rnaviralSPAdes, coronaSPAdes, and rnaSPAdes version 3.15.4
2. Trinity v2.13.2
3. MEGAHIT v1.2.9

### 2 Assembly command lines

#### 2.1 Quality trimming

Prior to assembly all reads underwent quality trimming using BBDuk:

```
bbduk.sh threads=16 trimpolya=15 qtrim=rl trimq=10 <input> <output>
```

#### 2.2 rnaviralSPAdes, coronaSPAdes, and rnaSPAdes

rnaviralSPAdes, and rnaSPAdes were run with the default parameters using 16 CPUs, e.g.:

```
rnaviralspades.py -t 16 -1 left.fastq -2 right.fastq
```

Noroviral HMMs were provided to `coronaspades.py` via `--custom_hmms` option:

```
coronaspades.py -t 16 -1 left.fastq -2 right.fastq --custom_hmms <hmms>
```

#### 2.3 Trinity

Trinity was run with the default parameters using 16 CPUs and 128 Gb maximum memory:

```
Trinity --seqType fq --left left.fastq --right right.fastq --CPU 16 --max_memory 128G
```

#### 2.4 MEGAHIT

MEGAHIT was run with the default parameters using 16 CPUs and utilizing 10% of available memory:

```
megahit -1 left.fastq -2 right.fastq -t 16 -m 0.1
```
